# Efficacy of PCV vaccine is primarily mediated by controlling pneumococcal colonisation density

**DOI:** 10.1101/547703

**Authors:** E.L. German, C. Solórzano, S. Sunny, F. Dunne, J.F. Gritzfeld, E. Mitsi, E. Nikolaou, A.D. Hyder-Wright, A.M. Collins, S.B. Gordon, D.M. Ferreir

## Abstract

Widespread use of Pneumococcal Conjugate Vaccines (PCV) has resulted in a reduction in nasopharyngeal colonisation and invasive pneumococcal disease caused by vaccine-types. In a double-blind, randomised controlled trial using the Experimental Human Pneumococcal Challenge (EHPC) model, PCV-13 (Prevenar-13) conferred 78% protection against colonisation acquisition and a reduction in bacterial intensity (AUC) in experimentally colonised volunteers as measured by classical culture. In this study, we used a multiplex quantitative PCR assay targeting *lytA* and pneumococcal serotype 6A/B *cpsA* genes to re-assess the experimental colonisation status of the same trial volunteers. Increase in detection of low-density colonised volunteers by this molecular method led to a decrease of PCV efficacy against colonisation acquisition (29%), as compared to classical culture (83%). For subjects who were colonised following pneumococcal challenge, PCV had a pronounced effect on decreasing colonisation density. These results have implications for vaccine efficacy and surveillance studies as they indicate that the success of PCV vaccination could primarily be mediated by the control of vaccine-type colonisation density which results in decreased transmission and the reported herd effect of PCVs. Studies assessing the impact of PCV should account for density measurements in their design.

Clinical trial registration with ISRCTN: 45340436

## Introduction

Pneumonia is a leading cause of death in children under 5 years worldwide, causing up to 1.4 million deaths annually [1]. Of these deaths, approximately 38% are caused by *Streptococcus pneumoniae* (pneumococcus) [2]. Current licensed pneumococcal conjugate vaccines (PCVs) are highly effective in protecting against invasive pneumococcal diseases caused by vaccine-type serotypes in children [3]. Immunization with PCVs also has beneficial indirect effects, conferring herd immunity to unvaccinated adults [4]. Disease and transmission prevention requires interruption of colonisation [3, 5]. Numerous studies have reported a reduction in vaccine-type pneumococcal colonisation acquisition and a concurrent decrease in colonisation density in individuals vaccinated with PCV, as measured by classical culture [6, 7].

We have previously reported the results of a double-blind, randomised controlled trial investigating the effect of the 13-valent PCV (PCV-13, Prevnar-13, Pfizer) on pneumococcal colonisation using the Experimental Human Pneumococcal Challenge (EHPC) model [8]. PCV showed 78% protection against acquisition of pneumococcus serotype 6B (BHN418) [9] (5/48 became colonised in PCV arm vs 23/48 in the control arm). Moreover, although the number of volunteers who became experimentally colonised following PCV vaccination was limited, we observed a significant reduction in nasal pneumococcal colonisation density (3 log difference in PCV arm compared to control arm at day 2). [8]

Interest in using molecular methods for colonisation detection is increasing because of their high sensitivity and, therefore, their ability to detect pneumococcus at low colonisation densities. Combining the results of classical culture and molecular methods has been suggested in order to improve accuracy of reported colonisation rates [10]. The *lytA* (autolysin) gene quantitative PCR (qPCR) strategy developed by the Centers for Disease Control (CDC) is currently the WHO-recommended culture-independent method to detect pneumococci [11, 12]. However, given the capacity of pneumococcus to exchange genes with other streptococci [13] a multiplex approach is valuable. Therefore, in this study, we employed a multiplex qPCR assay targeting *lytA* and pneumococcal serotype 6A/B *cpsA* genes to re-assess volunteer samples from our PCV study for experimental colonisation of 6B pneumococcus. We compared the results obtained by this molecular method with the results previously obtained by classical culture.

## Material and Methods

### PCV/EHPC Clinical trial

A trial investigating the efficacy of the 13-valent PCV vaccine against experimental human pneumococcal challenge was conducted in 2012. Study design and outcomes have been previously reported [8]. Briefly, 96 healthy volunteers aged 18-50 were vaccinated with either PCV-13 (PCV arm, n=48) or Hepatitis A vaccine (control arm, n=48). 4-5 weeks post-vaccination, volunteers were inoculated with 80,000 Colony-Forming Units (CFU) per nostril of live 6B pneumococcus (BHN418, sequence type 138) [8]. Nasal wash samples were taken and processed as described previously [14], before and after pneumococcal inoculation (at days 2, 4, 14 and 21). Samples were stored at −80°C. This trial was approved by The National Health Service Research and Ethics Committee (REC) (12/NW/0873 Liverpool) and was co-sponsored by the Liverpool School of Tropical Medicine and the Royal Liverpool and Broadgreen University Hospitals Trust. Informed consent was obtained from all volunteers.

### DNA extraction

300μl of D2, D7 and D14 nasal wash pellets was centrifuged for 7 minutes at 20,238xg. Following centrifugation, 300μl of lysis buffer with protease, 100μl of Zirconium beads (Stratech, Ely, UK) and 300μl of Phenol (Sigma-Aldrich, St Louis, MO, USA) was added to the pellet and the sample was disrupted twice for 3 minutes in a tissue homogenizer (Bertin Technologies, Montigny le Bretonneux, France) followed by 3 minutes on ice. The sample was centrifuged for 10 minutes at 9,391xg, and the upper aqueous phase was transferred to a tube pre-filled with 600μl binding buffer and 10μl magnetic beads. The samples were incubated at room temperature for 30-120 minutes then washed twice with 200μl of wash buffers 1 and 2. Magnetic beads were dried at 55°C for 10 minutes, eluted in 63μl of elution buffer and stored at −20°C. Lysis buffer, protease, binding buffer, magnetic beads, wash buffers 1 and 2, and elution buffer are part of the Agowa mag Mini DNA isolation kit (LGC Genomics, Berlin, Germany).

### Multiplex qPCR

We developed a novel multiplex qPCR based on methods previously published, using partial amplification of *lytA* [11] and 6A/B *cpsA* [15] genes. The sequences of the primers and probes used are: *lytA* forward primer: 5’-ACG-CAA-TCT-AGC-AGA-TGA-AGC-A-3’; *lytA* reverse primer 5’-TCG-TGC-GTT-TTA-ATT-CCA-GCT-3’; *lytA* probe: 5’-(FAM)-TGC-CGA-AAA-CGC-TTG-ATA-CAG-GGA-G-(BHQ-1)-3’; *cpsA* forward primer: 5’-AAG-TTT-GCA-CTA-GAG-TAT-GGG-AAG-GT-3’; *cpsA* reverse primer: 5’-ACA-TTA-TGT-CCA-TGT-CTT-CGA-TAC-AAG-3’; *cpsA* probe: 5’-(HEX)-TGT-TCT-GCC-CTG-AGC-AAC-TGG-(BHQ-1)-3’. The reaction mixture of 25 µl contained 0.6 µM of each *lytA* primer, 0.3 µM of *lytA* probe, 0.4 µM of each *cpsA* primer, 0.2 µM of *cpsA* probe, 12.5 µM of Taqman Gene Expression Master Mix (Applied Biosystems, USA) and 2.5 µL of extracted DNA. The qPCR reaction was run on a Mx3005P machine (Agilent Technologies, Santa Clara, CA, USA) on the following programme: 10 minutes at 95 °C followed by 40 cycles of 15 seconds at 95 °C and 1 minute at 60 °C. DNA from BHN418 serotype 6B, extracted using the QIAamp DNA mini kit (Qiagen, Hilden, Germany) and serially diluted 1:10 from 4.14×10^6^ copies in 2.5 µl, was used as a standard curve. A sample was considered positive if at least one duplicate had a CT value less than 40.

### Statistical Analysis

Risk Ratio, 95% confidence interval (CI) and p-value were calculated. Colonisation densities were determined from number of gene copies per well and analysed in GraphPad Prism v5 (GraphPad Inc.). Mean densities with standard deviations were also calculated.

## Results

### Colonisation acquisition rates by molecular methods

Samples from 90 of the 98 volunteers enrolled in the EHPC PCV study were available and included in these analyses. The re-assessment of the PCV/EHPC trial samples using the developed multiplex qPCR showed that all volunteers reported as colonisation-positive (acquired the inoculated bacterium) by classical culture were also found to be colonised with pneumococcus by molecular methods. However, the number of colonisation-positive volunteers increased in both PCV and control arms when using molecular methods. Positivity at any day for both *lytA* and *cpsA* was found in 22/45 (49%) volunteers in the PCV arm and 31/45 (69%) volunteers in the control arm (Table 1). The risk ratio by classical culture was 0.17 (95% CI, 0.07 to 0.46; P=0.0005) and by molecular methods was 0.71 (95% CI, 0.50-1.01; P=0.06). PCV conferred 83% protection against experimental pneumococcal colonisation by classical culture, and 29% protection by molecular methods.

**Table 1:** Comparison of numbers of colonised volunteers by detection method, study day and study arm

|  | Classical Culture |  | lytA/cpsA multiplex qPCR |  |
| --- | --- | --- | --- | --- |
|  | No. Colonised/Total No. (%) |  | No. Colonised/Total No. (%) |  |
|  | PCV | Control | PCV | Control |
| D2 | 2/38 (5) | 18/39 (46) | 11/38 (29) | 21/39 (54) |
| D7 | 4/43 (9) | 18/42 (43) | 16/43 (37) | 23/41 (56) |
| D14 | 1/9 (11) | 18/23 (79) | 2/9 (22) | 17/23 (74) |
| Any Day | 4/45 (9) | 23/45 (51) | 22/45 (49) | 31/45 (69) |

*Definition of abbreviations: PCV = pneumococcal conjugate vaccine; NW = nasal wash.*

When assessing the breakdown of colonisation-positive volunteers per study day, the percentage of experimentally colonised volunteers by *lytA*/*cpsA* multiplex qPCR was similar between D2 and D7 regardless of study arm (Table 1).

### Colonisation densities by molecular methods

Colonisation densities were lower in volunteers vaccinated with PCV compared to the control arm by both classical culture and molecular methods (Figure 1).

**Figure 1:**
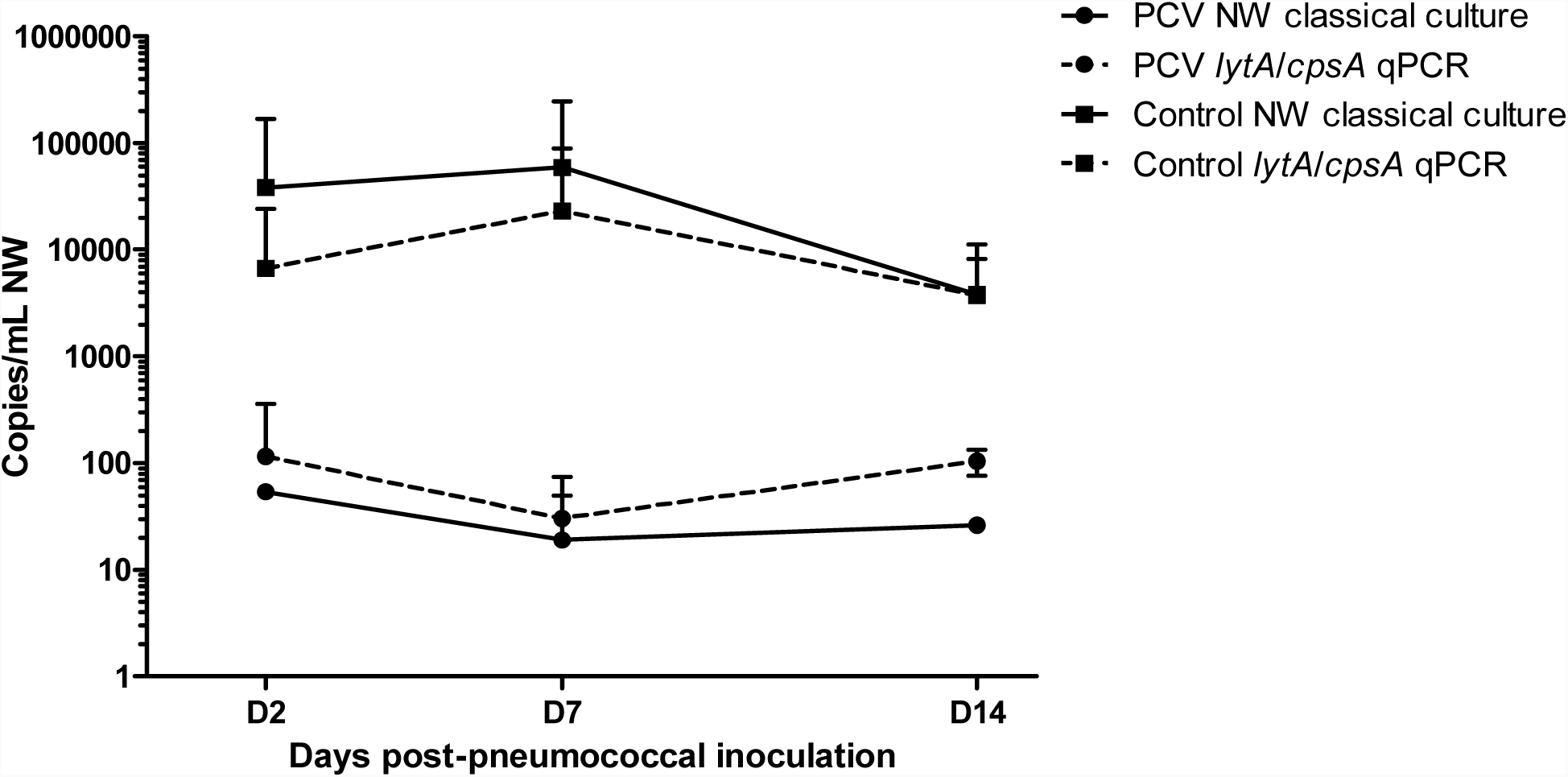
Colonisation densities by vaccination group, detection method and study time-point.

There was a significant (p<0.0001) correlation between densities calculated by classical culture and by *lytA*/*cpsA* qPCR for volunteers in both PCV and control arms (data not shown). 91% of samples positive by *lytA*/*cpsA* qPCR but not by classical culture had densities <30 DNA copies/ml. The mean colonisation density in the PCV arm by lytA/cpsA qPCR was 2.1 times higher than by classical culture at D2 and 1.6 times higher at D7. Conversely, the mean colonisation densities in the control arm for culture positive samples was 5.7 times lower by *lytA*/*cpsA* qPCR at D2 and 2.6 times lower by *lytA*/*cpsA* qPCR at D7 (Table 2).

**Table 2:** Comparison of colonisation density of colonised volunteers by detection method, study day and study arm

|  | Classical Culture |  | *lytA/cpsA* multiplex qPCR |  |
| --- | --- | --- | --- | --- |
| | CFU/ml Mean ± SD | | DNA copies/ml Mean ± SD | |
|  | PCV | Control | PCV | Control |
| D2 | 54 ± 2 | 38426 ± 130391 | 116 ± 244 | 6723 ± 17427 |
| D7 | 19 ± 31 | 59402 ± 187650 | 30 ± 44 | 23081 ± 66468 |
| D14 | 26 | 3823 ± 7300 | 105 ± 29 | 3740 ± 4452 |

*Definition of abbreviations: PCV = pneumococcal conjugate vaccine; CFU = colony forming units; SD = standard deviation; NW = nasal wash*

By either detection method, densities in the PCV arm were lower than in the control arm. The densities in the control arm decrease and those in the PCV arm slightly increase when measuring by multiplex qPCR.

## Discussion

In this study we re-assessed the experimental colonisation status of all volunteers from our PCV/EHPC trial using a multiplex qPCR. This qPCR targeted a specific region of *lytA* and serotype 6A/B *cspA* genes. Epidemiological studies have reported that the incidence of pneumococcal serotype 6A is very low in the UK [16]; this was further confirmed by screening 795 volunteers for the EHPC studies in Liverpool which found only one volunteer carrying a serotype 6A (Adler H et al., unpublished data). This gives reassurance that the combined detection of *lytA* and a *cspA* gene sequence common to serotypes 6A/B indicates the presence of the experimental pneumococcal 6B used in this study. We therefore report the results of our *lytA*/*cpsA* multiplex qPCR as the accurate numbers for molecular detection of experimental colonisation in these study participants.

The number of volunteers that became colonised after experimental pneumococcal inoculation by molecular methods was higher than the number previously reported by classical culture [8]. This increase in number is more pronounced in the PCV arm (23 vs 4 volunteers) than in the control arm (31 vs 23 volunteers), which translates into increased risk ratios, and therefore a decrease in the calculated protective efficacy of PCV against pneumococcal colonisation from 83% to 29%. Molecular methods, such as qPCR can detect DNA from dead bacteria. However, it is unlikely that this phenomenon had a major contribution to our findings. If this method were detecting remains of the pneumococcal inoculum administered on Day 0, we would expect to see higher colonisation rates at D2 than D7. However, the colonisation rates on both days are comparable.

Both detection methods demonstrate a lower colonisation density in volunteers vaccinated with PCV. The additional colonised volunteers detected in both arms by molecular methods are mostly colonised at a low density. Because volunteers vaccinated with PCV had already demonstrated low colonisation densities by classical culture, the addition of new colonisation-positive volunteers does not greatly alter the mean colonisation density. However, in the control arm, the additional low-density colonised volunteers have a larger effect in reducing the mean colonisation density. It has been suggested that colonisation density plays an underestimated but pivotal role in the development of pneumococcal disease and in transmission dynamics [17–19]. Our results further support the hypothesis that colonisation density is a determining factor for the clinical outcome and spread of *S. pneumoniae* and should be accounted for in the design of vaccine efficacy as well as colonisation surveillance studies.

## Conclusions

Using molecular methods, we have shown that PCV conferred 29% protection against experimental colonisation acquisition. This may indicate that the main protective mechanism of this vaccine is mediated by reduction of colonisation density, leading to a decreased risk of disease to vaccinated individuals as well as transmission resulting in the observed herd effects in vaccinated populations.

## Author Contributions

ADH-W, AMC, SBG and DMF designed and co-ordinated the PCV clinical trial; JFG, EM and DMF processed clinical samples; ELG, CS, SS, FD, JFG, EM and EN performed DNA extractions and qPCRs; ELG, CS, EM and EN developed the multiplex qPCR; ELG, CS and DMF analysed the data and drafted the manuscript. All authors have read and approved the manuscript.

## Acknowledgements

The authors would like to thank Dr Tao Chen for his advice on statistical analysis.

## Competing interests

Declarations of interest: none

This work was partially presented as a poster at the 10^th^ International Symposium on Pneumococci and Pneumococcal Diseases (ISPPD-10), 26^th^-30^th^ June 2016, abstract number 330.

